# Transcriptional, developmental, and functional parallels of lymphatic and venous smooth muscle

**DOI:** 10.1101/2024.07.18.604042

**Authors:** Guillermo Arroyo-Ataz, Alejandra Carrasco Yagüe, Julia C. Breda, Sarah A. Mazzilli, Dennis Jones

## Abstract

Lymphatic muscle cells (LMCs) are indispensable for lymphatic vessel contraction and their aberrant recruitment or absence is associated with both primary and secondary lymphedema. Despite their critical role in lymphatic vessel function, the transcriptomic and developmental basis that confer the unique contractile properties to LMCs are largely undefined. In this study, we employed single-cell RNA sequencing (scRNAseq), lineage tracing and *in vivo* imaging to investigate the basis for the hybrid cardiomyocyte and blood vascular smooth muscle cell (SMC) characteristics that have been described for LMCs. Using scRNAseq, the transcriptomes of LMC and venous SMCs from the murine hindlimb exhibited more similarities than differences, although both were markedly distinct from that of arteriole SMCs in the same tissue. Functionally, both lymphatic vessels and blood vessels in the murine hindlimb displayed pulsatile contractility. However, despite expressing genes that overlap with the venous SMC transcriptome, through lineage tracing we show that LMCs do not originate from Myh11+ SMC progenitors. Previous studies have shown that LMCs express cardiac-related genes, whereas in our study we found that arteriole SMCs, but not LMCs, expressed cardiac-related genes. Through lineage tracing, we demonstrate that a subpopulation of LMCs and SMCs originate from WT1+ mesodermal progenitors, which are known to give rise to SMCs. LMCs, however, do not derive from Nkx2.5+ cardiomyocyte progenitors. Overall, our findings suggest that venous SMCs and LMCs and may derive from a related mesodermal progenitor and adopt a similar gene expression program that enable their contractile properties.

## Introduction

Lymphatic muscle cells (LMCs) embedded in the wall of collecting lymphatic vessels contract to facilitate lymphatic pumping, which is a major driver for the removal of fluid from tissues^1^. Several studies have revealed the importance of LMCs in health and disease, providing insights into how LMC dysfunction contributes to primary and secondary lymphedema, and various chronic inflammatory diseases, including obesity, arthritis, inflammatory bowel disease, and aging^2–7^. Our previous work demonstrated that disorganized and incomplete LMC coverage of collecting lymphatic vessels following *Staphylococcus aureus* infection was associated with permanent impairment of collecting lymphatic vessel contraction and lymph flow^8^.

Developing targeted treatments, potentially involving pharmacological targets or regenerative therapies, that can modulate LMC function and restore lymphatic contractility may represent a promising approach to address deficiencies in lymphatic pumping. Yet, our ability to modulate LMC function is hindered by our incomplete understanding of their molecular signature, impeding precise therapeutic interventions targeting LMCs.

The limited characterization of LMCs, primarily made through the analysis of bulk cells or functional interventions in isolated vessels, has revealed a distinctive hybrid nature, sharing features with both smooth and striated muscle types^7^, including cardiac muscle and skeletal muscle. Indeed, LMCs in different vascular beds reliably express canonical smooth muscle cell (SMC) markers including alpha smooth muscle actin (α-SMA), smooth muscle myosin heavy chain (SM-MHC, Myh11), integrin-α8, PDGFR-β, collagen IV, and desmin^9–14^. Moreover, similar expression patterns of L-type calcium channels in LMCs and regulation of calcium signaling through myosin light chain kinase indicate a common functional profile with SMCs^15–17^. However, little evidence exists to confirm the hybrid features of LMCs. Lineage tracing using Pax7 and Myod recently revealed that LMCs do not develop from skeletal muscle tissue^18^. Apart from the reported expression of CaV 3.1 and CaV 3.2 T-type calcium channels, which are also expressed in cardiac tissue, there is limited evidence indicating an enrichment of cardiac-related genes in LMCs ^15,19,20^.

Considering the hybrid features of LMCs, it remains largely unclear whether LMCs arise from distinct precursor cells, leading to diversity among LMCs, or whether they originate from a common progenitor and develop a unique expression profile. On the other hand, it is well documented that SMCs originate from diverse progenitor sources, with prominent contributions from mesodermal and neural crest cells^21–25^. Whether cardiac precursors contribute to LMCs and confer cardiac-like properties to these cells has not been established.

Here, we used lineage-tracing, single-cell RNA sequencing (scRNAseq), and intravital imaging to elucidate the transcriptional characteristics, developmental origins, and contractile properties of both LMCs and SMCs in the same anatomical site in mice. scRNAseq analysis revealed a high degree of transcriptional similarity between LMCs and venous SMCs, while both cell types exhibited contractility *in vivo*. In contrast to previous studies, we found that arterial SMCs near lymphatic vessels, but not LMCs, expressed cardiomyocyte markers. Additionally, lineage tracing reveals that LMCs do not originate from SMCs. However, both LMCs and SMCs may emerge from clonally related progenitor cells that do not include cardiomyocyte progenitors.

## Methods

### Mice and treatments

All procedures were performed according to the guidelines of the Institutional Animal Care and Use Committee of Boston University.

<u>For assessing embryonic LMC recruitment to lymphatic vessels</u>, we used wild type C57BL/6J mice (Stock number 000664, The Jackson Laboratory). Pregnant dams were anesthetized with ketamine/xylazine (K/X, 0.2 mL/20g) and euthanized by cervical dislocation to extract embryos at days 15.5, 17.0 or 17.5 days post-coitum. Neonatal mice (P0 and P7) were euthanized by decapitation. Popliteal vessels and mesenteries were harvested from the embryos and pups.

<u>For conventional fate mapping</u> of WT1 and Nkx2.5 progenitors, we used Nkx2.5^IRESCre^ [B6.129S1(SJL)-*Nkx2-5^tm2(cre)Rph^*/J, Stock number 024637, The Jackson Laboratory]^26^ and WT1^GFPCre^ [STOCK *Wt1^tm1(EGFP/cre)Wtp^*/J, Stock number 010911, The Jackson Laboratory]^27^ crossed with Ai9-tdT [B6.Cg-*Gt(ROSA)26Sor^tm9(CAG–tdTomato)Hze^*/J, Stock number 007909, The Jackson Laboratory]^3^ mice. Ai9 mice have a floxed STOP cassette upstream of tdTomato, and when bred to mice that express Cre recombinase, the resulting offspring presents expression of tdTomato in all the cells that express Cre, as well as their progeny. Popliteal vessels were harvested at P7 and >3 weeks. Mesenteries were harvested at P7.

<u>For conditional fate mapping</u> of WT1 and Myh11 progenitors, we used WT1^CreERT^^2^ [STOCK *Wt1^tm2(cre/ERT2)Wtp^*/J, Stock number 010912, The Jackson Laboratory]^27^ and SMMHC-iCreER^T2^ [B6.FVB-Tg (Myh11-icre/ERT2)1Soff/J, Stock number 019079, The Jackson Laboratory]^28^ crossed with Ai9-tdT mice. To induce translocation of CreERT2 in conditional Myh11xAi9 embryos, pregnant dams were orally administered one dose of tamoxifen (75 mg/kg body weight in 100 µL of corn oil) 14.0 days post-coitum. For conditional WT1xAi9 fate mapping, postnatally induced pups were injected intraperitoneally (50 µL of 2 mg/ml) at day 4 and 7 after birth. Popliteal vessels, mesenteries and kidneys were harvested from mice induced at postnatal day 11 (P11). For induction of adult Myh11xAi9 animals, one oral dose of tamoxifen was used (75 mg/kg body weight in 100 µL of corn oil), and the popliteal vessels were harvested 4 days after.

<u>For WT1 specific ablation</u>, we crossed WT1^GFPCre^ mice with ROSA26iDTR [C57BL/6-*Gt (ROSA)26Sor^tm1(HBEGF)Awai^*/J, Stock number 007900, The Jackson Laboratory]^29^ mice. To induce ablation in WT1xDTR mice, adults were anesthetized with K/X (0.2 mL/20g) and administered subcutaneous injections of diphtheria toxin (5 ng/50 µl) in the hindlimb (parallel to the saphenous vein) for three consecutive days. Hindlimbs were collected four days after the final diphtheria toxin injection (day 7).

### Harvesting of vessels and organs

Adult popliteal lymphatic vessels (popliteal lymphatic vessels) and popliteal blood vessels (saphenous vein) were harvested following a previously described procedure^30^. Briefly, mice were anesthetized with K/X (0.2 mL/20g) and 4% Evans Blue dye was subcutaneously injected into the footpad to highlight the popliteal lymphatic vessels. Next, hindlimb skin was removed to expose the popliteal vessels. Once exposed, the popliteal lymphatic vessels and the saphenous vein were dissected out of the surrounding fat and underlying muscles using a stereomicroscope, followed by a PBS wash. For neonatal and embryonic tissues, the animals were euthanized by decapitation and the hindlimbs were isolated. Then, the skin of the entire limb was carefully removed using a stereomicroscope, trimmed, and processed whole till imaging.

Mesenteries were harvested from embryos and neonates following a modified version of a previously described protocol^31^. Briefly, animals were placed on their back and an incision was made on the midsection of their abdomen to expose the peritoneal cavity. Once the large intestine was identified, it was carefully pulled out of the peritoneal cavity paying attention not to tear the mesothelial membrane. Finally, the intestine and attached structures were washed in PBS, and with help of a stereomicroscope the mesentery was separated from the intestine to continue processing.

Heart of some adult and neonatal animals were harvested as positive controls for Pln staining. To this purpose, we opened the chest and abdominal cavities as described by others and harvested the organs of interest^32^.

### Immunofluorescent whole-mount staining

For whole-mount staining of popliteal vessels^8^, extracted tissues were fixed in 4% paraformaldehyde for 1-4 hours at room temperature, washed in PBS and permeated with 0.5% TritonX-100 (T-X) in PBS O/N at 4°C. Of note, for fixation-sensitive antigens, fixation was shortened to 15 minutes. The tissues were then washed 3 x 20 min with PBS-glycine wash buffer (0.2 M glycine) at room temperature and blocked for 1 hour with 10% normal donkey serum at room temperature. Next, the samples were incubated for 48-72 hours with primary antibodies at 4°C, washed with PBS-glycine wash buffer 3 x 20 min, and incubated overnight with secondary antibodies at 4°C. The samples were then washed with PBS and mounted for imaging.

For whole mount staining of mesenteric vessels^10^, extracted tissues were fixed in 4% paraformaldehyde for 6 hours at 4°C and washed in PBS. The tissues were then blocked and permeated with PBS 0.3% triton-X 100 (PBSTx) with 3% BSA for 3 h at room temperature. Next, tissues were incubated overnight at 4°C with primary antibodies, washed with 0.3% triton-X 100 3×1h at room temperature on a rocking table, and incubated overnight with secondary antibodies at 4°C. The samples were then washed with PBS and mounted for imaging with ProLong™ Gold Antifade Mountant with DAPI (ThermoFisher, P36931).

Primary antibodies included mouse anti-α-SMA, Alexa Fluor^TM^ 488 (1:200; 53976082, Invitrogen), mouse anti-α-SMA (1:200; 14976082, Invitrogen), rabbit anti-Pln (1:100; A23750, Abclonal), rabbit anti-DOG-1 (1:100; MA5-16358, ThermoFisher), mouse anti-Enpp2 (1:100; ab77104, Abcam), rabbit anti-AEBP1 (1:100; gift from Dr. Matthew Layne) and goat anti-Prox1 (1:40; AF2727, R&D Systems). Where applicable, we used secondary antibodies conjugated with Alexa Fluor^TM^ 488, 647, or Cy3 fluorophores from Jackson ImmunoResearch Laboratories (1:100 for staining of all tissues). Samples were imaged using a Leica SP5 microscope with Leica Application Suite software. Images were acquired at 10X, 20X, 40X or 60X using sequential scanning mode. Area quantification and cell counting were performed using ImageJ. Intensity was enhanced for figures 2F (Pln panels), 3A (lower panels), 4B (central panel), 4D (all panels), 5C (all panels), 5E (all panels), 6C (Upper Prox1 panel), 7E (Upper Prox1 panel) and S4C (tdT panel). Background was subtracted from 3D projections of 2H (Pln panel) and 7E (Upper Prox1 panel).

### scRNAseq

To obtain single cell suspensions from popliteal lymphatic vessels and blood vessels, adult male and female WT1xAi9 mice were anesthetized with K/X (0.2 mL/20g) and administered 4% Evans blue dye footpad injections. Popliteal vessels were then exposed, and we used a previously described method^30^ to generate single cell suspensions of the vessels. Briefly, minced vessels were incubated for 60 minutes at 37°C with agitation in an enzymatic cocktail consisting of 1 mg/ml collagenase I (Worthington, LS004196), 1 mg/ml Dispase II (Gibco, 17105041). The resulting cell suspension was then filtered through a 70 µm nylon filter (Fisher Scientific, 22363548) and digestion enzymes were inactivated by adding FBS. In total, we performed this protocol on three sets of two mice each. For the first set, we extracted the blood vessels and popliteal lymphatic vessels and hashed the samples with TotalSeq™-A0306 anti-mouse Hashtag 6 Antibody (Biolegend, 155811) and TotalSeq™-A0305 anti-mouse Hashtag 5 Antibody (Biolegend, 155809) according to the manufacturer’s instructions. Only hashed cells from blood vessels are presented. For the other two sets we extracted the popliteal lymphatic vessels and performed the digestion for 30 minutes and adding DNAse (Milipore Sigma, 04716728001) and Liberase (Milipore Sigma, 05401020001) to the enzymatic cocktail. For scRNAseq, single-cell suspensions were dispensed onto a Chromium Instrument (10x Genomics) and targeted capture of 10,000 cells per sample. Our scRNAseq library was made using the Next GEM Single Cell 3’ Reagent Kit (10X Genomics). cDNA libraries were combined and loaded in a flow-cell for sequencing using a NextSeq2000 (Illumina) to obtain a sequencing depth of at least 50,000 reads per cell.

### scRNAseq data processing

Creation of FASTQ files were performed using BaseSpace. For hashed samples, cells were demultiplexed using the HTODemux function in Seurat, tdTomato sequence was incorporated into the mouse genome reference (ENSEMBL, GRCm38) and single-cell sequencing reads subjected to preprocessing using Cell Ranger analysis pipeline. Expression matrices were imported into R and analyzed using the Seurat Package^33^. First, low quality cells (≥ 30,000 counts library size, ≤ 200 expressed genes, > 10% mitochondrial genes) were removed from the data set. After quality control, there were 2376 popliteal lymphatic vessel-derived cells and 856 blood vessel-derived cells from n=6 mice. To visualize the data, dimensionality reduction was performed by Principal Component Analysis (PCA), Uniform Manifold Approximation and Projection (UMAP), and Harmony using the Seurat functions RunPCA, RunUMAP, and RunHarmony. Clusters that formed the data were defined using the functions FindNeighbors (dims 1:30) and FindClusters (resolution=0.8). The markers of each cluster were found using the functions FindMarkers and FindAllmarkers, and those that shared cell identities (except for SMCs and LMCs) were combined. Unbiased analysis was also performed with the function using the EnrichR data base using the function DEenrichRPlot to confirm cell identities.

Raw scRNA sequencing data of popliteal lymphatics and saphenous vein cells have been deposited in the Gene Expression Omnibus (GEO) GSE252376.

#### Quantitative real-time Polymerase Chain Reaction (qPCR)

RNA was isolated using a RNeasy mini kit and QIAshredder (Qiagen, Germantown, MD) to homogenize the cells. The quality and quantity of RNA were determined spectrophotometrically at 260 and 280 nm, respectively, using a NanoDrop ND-1000 spectrophotometer. Purified mRNA was reverse transcribed to cDNA using the Verso cDNA Synthesis Kit according to the manufacturer’s (ThermoScientific) instructions. Next, real-time PCR was performed according to the manufacturer’s instructions using iTAQ Universal SYBR Green Supermix (Bio-Rad (Hercules, CA, cat # 172-5124) on a Bio-Rad CFX Connect Real-Time PCR Detection System to amplify the following murine genes: *Enpp2*, (forward) 5’-GGATTACAGCCACCAAGCAAGG-3’, (reverse) 5’-ATCAGGCTGCTCGGAGTAGAAG3’; *Aebp1* (forward) 5’-GAGGTGGTAACTACTGACAGCC -3’, (reverse) 5’-CCAGGCTGTATGTGCGAGTGAT’; *Pcdh7* (forward) 5’-CAGATCACTGCTCCGAGTACAG -3’, (reverse) 5’-CCGCTGTCATAGCAACTCTGC A-3’

### Measuring lymphatic vessel contraction

To characterize venous and lymphatic contractility, contraction studies were performed and analyzed similarly as previously described^8,34,35^. Briefly, mice (n=3) between 5-6 weeks were anesthetized with ketamine (100mg/kg) and xylazine (10mg/kg)) and given a dorsal footpad injection of 2 µL 2% FITC-Dextran in sterile saline. Then, the popliteal vasculature was exposed by excising the dorsal hindlimb skin and connective tissue near the popliteal lymphatic vessels. The dye-filled popliteal lymphatic vessels are then imaged using an inverted fluorescence microscope (EVOS™ M5000 Imaging System). Each image capture sequence is conducted within the living animal over 90 seconds (30 individual images separated by 3 seconds). The areas of the popliteal lymphatic vessel and saphenous vein was measured using ImageJ of all images and plotted over time. To calculate the frequency, every peak in this plot was considered a complete cycle.

### Primers for genotyping

WT1^GFPCre^:

Control primer Forward 5’CCT ACC ATC CGC AAC CAA G 3’
Control primer Reverse 5’CCC TGT CCG CTA CTT TCA GA 3’
WT1 primer Forward 5’ATC GCA GGA GCG GAG AAC 3’
WT1 primer Reverse 5’GAA CTT CAG GGT CAG CTT GC 3’

WT1^CreERT2^:

Common primer Forward 5’ATC GCA GGA GCG GAG AAC 3’
Control primer Reverse 5’GAA GGG TCC GTA GCG ACA 3’
WT1 primer Reverse 5’GCA AAC GGA CAG AAG CAT TT 3’

Nkx2.5^IRESCre^:

Nkx2.5 primer Forward 5’ TTA CGG CGC TAA GGA TGA CT 3’
Control primer Forward 5’ GAG CCT GGT AGG GAA AGA GC 3’
Common primer Reverse 5’ GTG TGG AAT CCG TCG AAA GT 3’

For WT1xDTR mice, WT1^GFPCre^ primers were used to confirm WT1 transgene, and DTR was homozygous.

### Statistics

Statistical analysis was performed using SPSS. Unpaired t tests were used to compare 2 data sets, and ANOVA and post-hoc tests (Games-Howell, Tukey’s test) were used to compare multiple data sets.

## Results

### Transcriptional profiling of lymphatic vessels and blood vessels from murine hindlimb

To transcriptionally profile LMCs and SMCs and potentially identify genes distinguishing between the two, we employed single cell RNA-sequencing (scRNAseq). We isolated lymphatic vessels afferent to the popliteal lymph node (popliteal lymphatic vessels) and adjacent superficial saphenous veins from the hindlimbs of mice using a surgical technique that we previously described^30^. Subsequently, single cells from respective samples underwent independent library preparation and sequencing using the 10x Genomics platform. Once the popliteal lymphatic vessel and saphenous vein libraries were prepared, we aligned them to a reference genome using Cell Ranger. Finally, by using Seurat, we identified 2,376 high quality single cells from popliteal lymphatic vessels and 856 single cells from saphenous vein. Uniform manifold approximation and projection (UMAP) for dimensionality reduction revealed 16 distinct clusters within the integrated popliteal lymphatic vessel and blood vessel samples, and cellular identities for these clusters were determined based on the enrichment of well-established lineage markers^36–47^ (Fig. 1A, B). Based on these analyses, we identified LMCs, arteriolar and venous SMCs (aSMCs and veSMCs respectively), endothelial cells (ECs), fibroblasts, weakly contractile perivascular cells (PCs), Schwann cells (SCs), macrophages, T-cells, B-cells, granulocytes, platelets and erythroid cells in both popliteal lymphatic vessel and saphenous vein derived data sets (Fig. 1B, C, S1A). Notably, despite originating from different vessels (Fig. 1C), we found that LMCs (190) and veSMCs (231) group together (veSMC/LMC cluster), suggesting a similar transcriptional identity.

**Figure 1.**
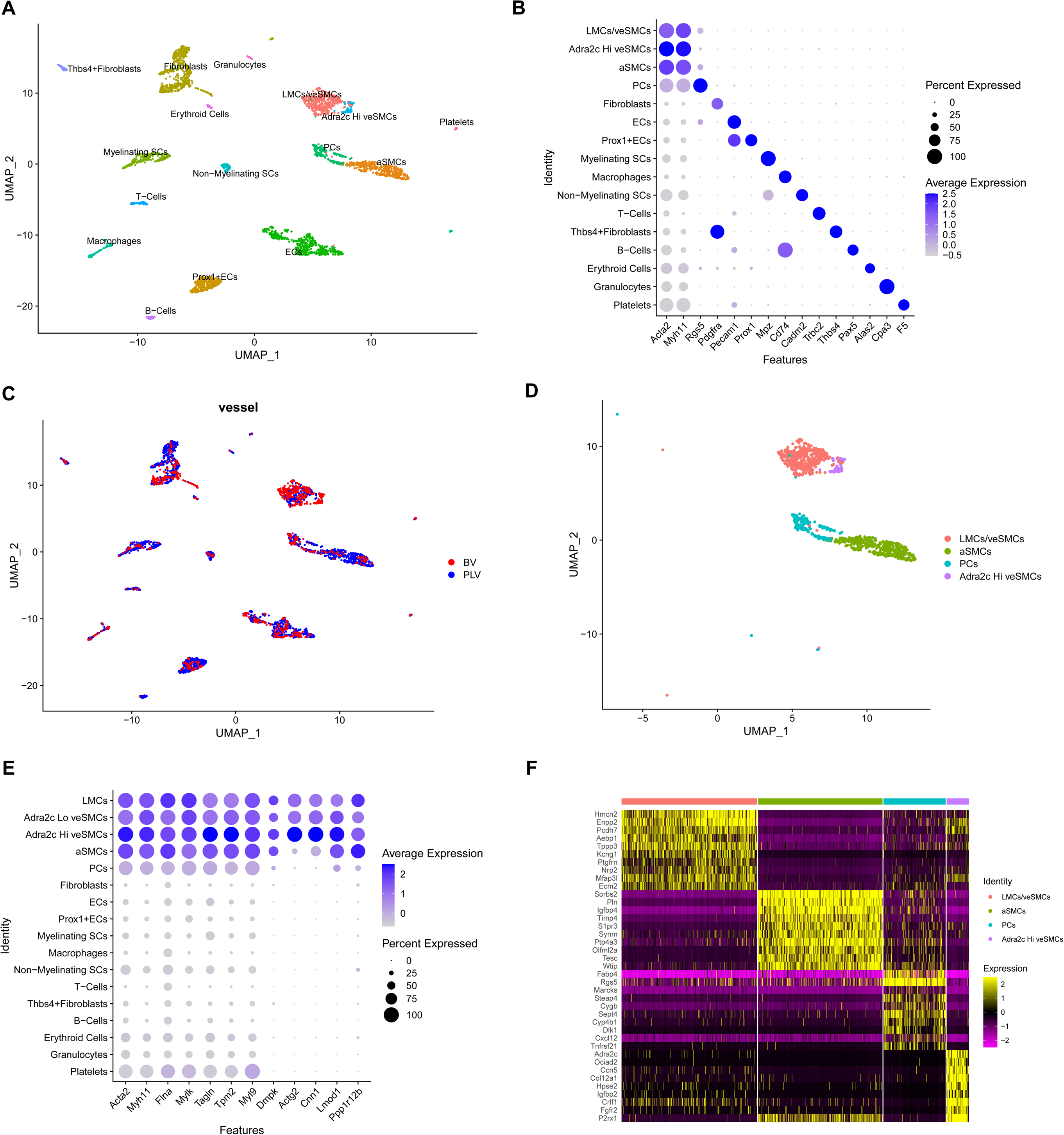
Lymphatic muscle cells show transcriptional similarities with venous smooth muscle cells. **A.** UMAP of 3232 single cells from merged dataset containing cells from popliteal lymphatic vessels (PLV) and saphenous vein (BV) of WT1 x Ai9 mice. Major cell types are color coded. **B.** Dot plot showing marker genes for clusters in A. **C.** UMAP color coded to show PLV or BV origin of cells. **D.** Color coded UMAP of contractile clusters from murine popliteal lymphatic vessels and saphenous vein. **E.** Dot plot with relative expression of selected canonical markers for contractile smooth muscle cells. **F.** Heatmap of the top differentially expressed genes (p-value) between each of the contractile clusters.

### scRNAseq of popliteal vascular branch reveals heterogeneity of contractile cell populations

In total, four contractile subpopulations from popliteal lymphatic vessels and saphenous vein were identified based on α-SMA (encoded by *Acta2*) and Myh11 expression (Fig. 1B, D). Recently, a panel of genes (*Acta2, Myh11, Tagln, Cnn1, Lmod1, Actg2, Tpm2, Ppp1r12b, Myl9, Dmpk, Mylk* and *Flna*) were shown to be conserved between SMCs across different organs^48,49^. All genes in this panel were enriched in three of the four contractile populations (Fig. 1E).

LMCs clustered together with veSMCs (Adra2c^lo^ veSMCs) in the LMC/veSMC cluster and accordingly, were enriched for many shared genes (Fig. 1F). The most markedly enriched genes of the cluster containing LMCs and veSMCs, based on p-value, were Ectonucleotide Pyrophosphatase/Phosphodiesterase 2 (*Enpp2*)/Autotaxin and the extracellular matrix protein Hemicentin 2 (*Hmcn2*). Enpp2 has a role in blood vessel stabilization and produces most of lysophosphatidic acid in serum, which facilitates angiogenesis and SMC proliferation^50–52^. We also identified a smaller subpopulation of veSMCs (Adra2c^hi^ veSMCs) that clustered separately from the major veSMC population and was enriched in Adrenoreceptor alpha-2c (*Adra2c*), an adrenergic receptor (Fig. 1F, S1B). Adra2c mediates neurotransmitter effects and regulates blood pressure through vascular smooth muscle contraction^53^. In addition to *Adra2c* expression, the *Adra2c* enriched veSMC cluster was also highly enriched in *P2rx1*, which encodes the purinergic receptor P2X1. This G-coupled protein receptor is associated with SMC contraction triggered by neuronal stimulation^54^.

A direct comparison between LMCs and veSMCs in the LMC/veSMC revealed genes that were differentially expressed between the cell types (Fig. S1C). *Mcam, Itga1, Nr2f2,* and *Cav1*, genes associated with signal transduction, were enriched in LMCs^55,56^ while *Col3a1* and *Sparc*, an extracellular matrix (ECM) and ECM-associated protein, respectively, were increased in veSMCs^56^. Moreover, Enrichr analysis indicated an upregulation in genes associated with ECM production and organization in veSMCs (Fig. S1D). In summary, unsupervised clustering reveals that LMCs and veSMCs group together, owing to the similarity in their gene expression profiles. However, variations in gene expression profiles were identified between the two cell types.

The second largest contractile population was comprised of cells enriched in Phospholamban (*Pln*, S1B). Cells in this cluster were also highly enriched genes in *Sorbs2*, *Igfbp4, Cspg4,* and *Rgs4* (Fig. 1F, S1E). Interestingly, Pln and sorbin and SH3 domain containing protein 2 (Sorbs2) are intricately linked to calcium transport and structural integrity in cardiomyocytes, respectively^57,58^. Pln, Rgs4 and neuron-glial antigen 2 (NG2), the protein encoded by *Cspg4*, are also expressed in vascular smooth muscle^48,59,60^. However, lineage tracing studies have shown that NG2 is not expressed in adult LMCs or veSMCs, and that this protein is primarily expressed by arteries and arterioles^18,48,61^. Additionally, while LMCs and veSMCs were highly enriched in Neuropilin2 (*Nrp2*), a marker of venous and lymphatic vessels^62–66^, cells within the *Pln* enriched cluster did not show any *Nrp2* expression (Fig. 1F). This suggested that LMCs were not the only contractile cell type contained within the single cell suspension derived from popliteal lymphatic vessels (Fig. 1C). Thus, based on their arteriole gene signature^48^, we refer to the cluster of Pln+ cells as arteriole SMCs (aSMCs).

A small subset of contractile cells was enriched for the mural cell markers regulator of G protein signaling 5 (Rgs5)^67^ and fatty acid binding protein 4 (Fabp4)^68^. Relative to the other contractile populations, these cells showed low expression of *Acta2*, *Myh11*, *Mylk* and *Tagln, Flna, Tpm2, Myl9*, and negligible expression of other signature SMC genes. We refer to these mural cells as weakly contractile perivascular cells (PCs, Fig. 1E).

For all contractile populations, we further analyzed the expression of Ca^2+^(*Cacna1c*, *Cacna1g*, *Cacna1h*), Na^+^ (*Scn3a*) and K^+^ channels (*Kcnj8* and *Kcnma1*), previously reported to be expressed in lymphatic vessels (Fig. 2A)^15,16,69–71^. *Cacna1c,* which encodes the Cav1.2 voltage gated L-type calcium channel, was enriched in LMCs and veSMCs, although its expression was maintained across the other contractile clusters. Similarly, *Scn3a* and *Kcnma1* were strongly expressed in LMCs and showed variable expression in other contractile clusters. In contrast, the T-type Ca^2+^ channels CaV3.1, CaV3.2 (*Cacna1g*, *Cacna1h*) were exclusively expressed in the aSMCs. Other genes previously shown to be expressed by LMCs, including *Prrx1, Itga8, Des* and *Ano1*, were also highly enriched in both LMCs and SMCs (Fig. 2A)^12,13,18,72^. Anoctamin 1 (Ano1), a calcium activated chloride channel, is critical for regulation of membrane potential and contractile responses in blood vessels and lymphatic vessels^72^. Since both LMCs and veSMCs exhibited similar RNA levels of *Ano1*, we validated protein expression of this marker in popliteal lymphatic vessels and saphenous veins by whole-mount immunofluorescence (Fig. 2B). Next, we sought to confirm the expression of additional genes enriched in both LMCs and veSMCs (Fig. 1F, 2C). To this end, popliteal lymphatic vessels and saphenous veins were isolated and quantitative PCR was performed on respective vessels to detect *Enpp2*, *Pcdh7*, and *Aebp1*. Transcript levels of indicated genes were similar between popliteal lymphatic vessels and saphenous veins (Fig. 2D), corroborating the findings of single-cell RNA sequencing. Whole mount immunofluorescent staining confirmed the expression of Enpp2 and Aebp1 in both collecting lymphatic vessels and veins (Fig. 2E, F).

**Figure 2.**
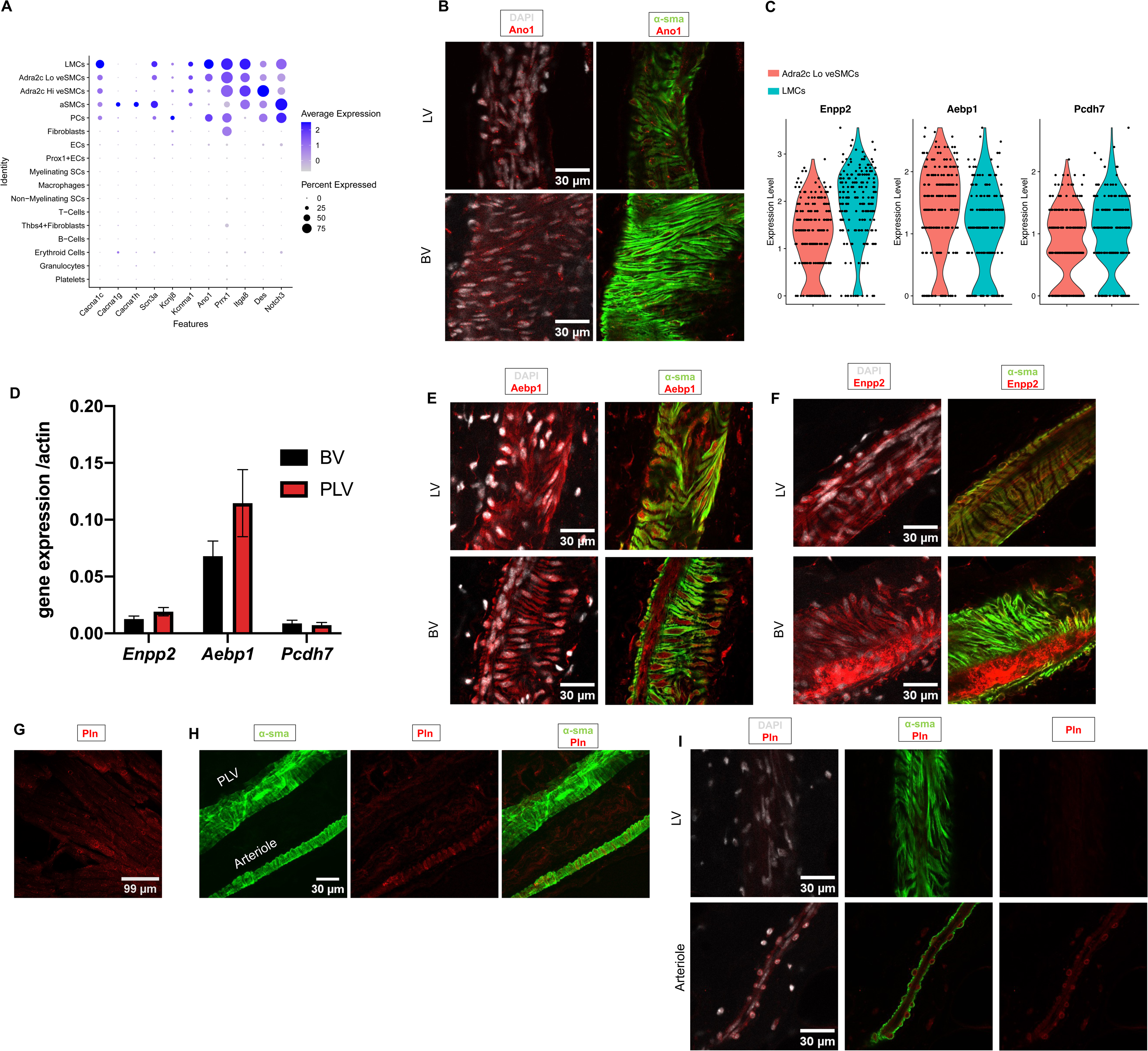
Validation of contractile populations in murine collecting lymphatic vessels. **A.** Dot plot showing relative expression of selected lymphatic markers. **B.** Representative 3D projection of Ano1 immunostained whole mount popliteal lymphatic vessels (PLV) and saphenous vein (BV). Ano1 (red) is overlaid with DAPI (white) on the left panel. Ano1 is overlaid with α-smooth muscle actin (sma), green, on the right panel. Original magnification 40X; Scale bar 30 µm, **C.** Violin plot of select top genes from the LMC/veSMC cluster (Fig. 1F). **D.** PLV and BVs were isolated to measure relative gene expression of select top genes displayed in Fig. 2c. Gene expression calculated using 2ΔCT method, as normalized to β-actin (n=3 biological replicates). Representative 3D projection of whole mount immunostained inguinal-axillary lymphatic vessel (LV) and adjacent thoracoepigastric vein (BV) with Aebp1 (**E**) and Enpp2 (**F**). Overlay with DAPI (white, left column) and α-sma (green, right column) is shown. Original magnification 40X; Scale bar 30 µm **G.** 3D projection of Pln immunostained adult heart (Pln positive control). Original magnification 40X; Scale bar 99 um. **H.** 3D projection of Pln (red) immunostained popliteal lymphatic vessel (PLV) and adjacent arteriole. α-sma (green). Original magnification 40X; Scale bar 30 µm. **I.** 3D projection of Pln (red) immunostained inguinal-axillary lymphatic vessels (LV, top panel) and adjacent arteriole (bottom panel. Overlay with DAPI (white) and α-sma (green) is shown. Original magnification 40X; Scale bar 30 µm.

We next performed whole-mount staining of the popliteal vascular branch using an anti-phospholamban antibody. As expected, phospholamban was detected in the sarcoplasmic reticulum of cardiomyocytes, which were stained as a positive control (Fig 2G). In both the popliteal vascular branch and inguinal axillary vascular branch, phospholamban staining localized to smooth muscle cells of arterioles, but not lymphatic vessels (Fig. 2H, I).

### The saphenous vein in mice exhibits phasic contractions

Since the transcriptional profiles of LMCs and saphenous vein SMCs showed substantial overlap, we next asked whether they had similar contractile properties. The phasic contractility of LMCs has been extensively characterized in the literature^7^, but has rarely been studied for venous SMCs. However, prior studies demonstrate that specific veins, such as the portal vein, can exhibit phasic contractility^49^ and consequently we hypothesized that saphenous veins were capable of this type of contraction. To test this hypothesis, we performed intravital imaging of popliteal lymphatic vessels and saphenous veins to determine the variation of their area over time. FITC-dextran fluorescence was used to visualize lymphatic vessels, while the contrast from red blood cells delineated blood vessels. Both vessels displayed diastolic and systolic phases, confirming phasic contraction in each (Fig. 3A, B). Notably, the contraction frequency in saphenous veins was significantly faster than that of popliteal lymphatic vessels (7.11 vs 3.33 cycles/min, p=0.005; Fig. 3C), while popliteal lymphatic vessels showed a higher variation between their maximum and minimum area (1.16 vs 1.65, p=0.0041; Fig. 3D). These data suggests that LMCs and saphenous vein SMCs have contractile properties, although the rate and force of contraction show variation between the vessels.

**Figure 3.**
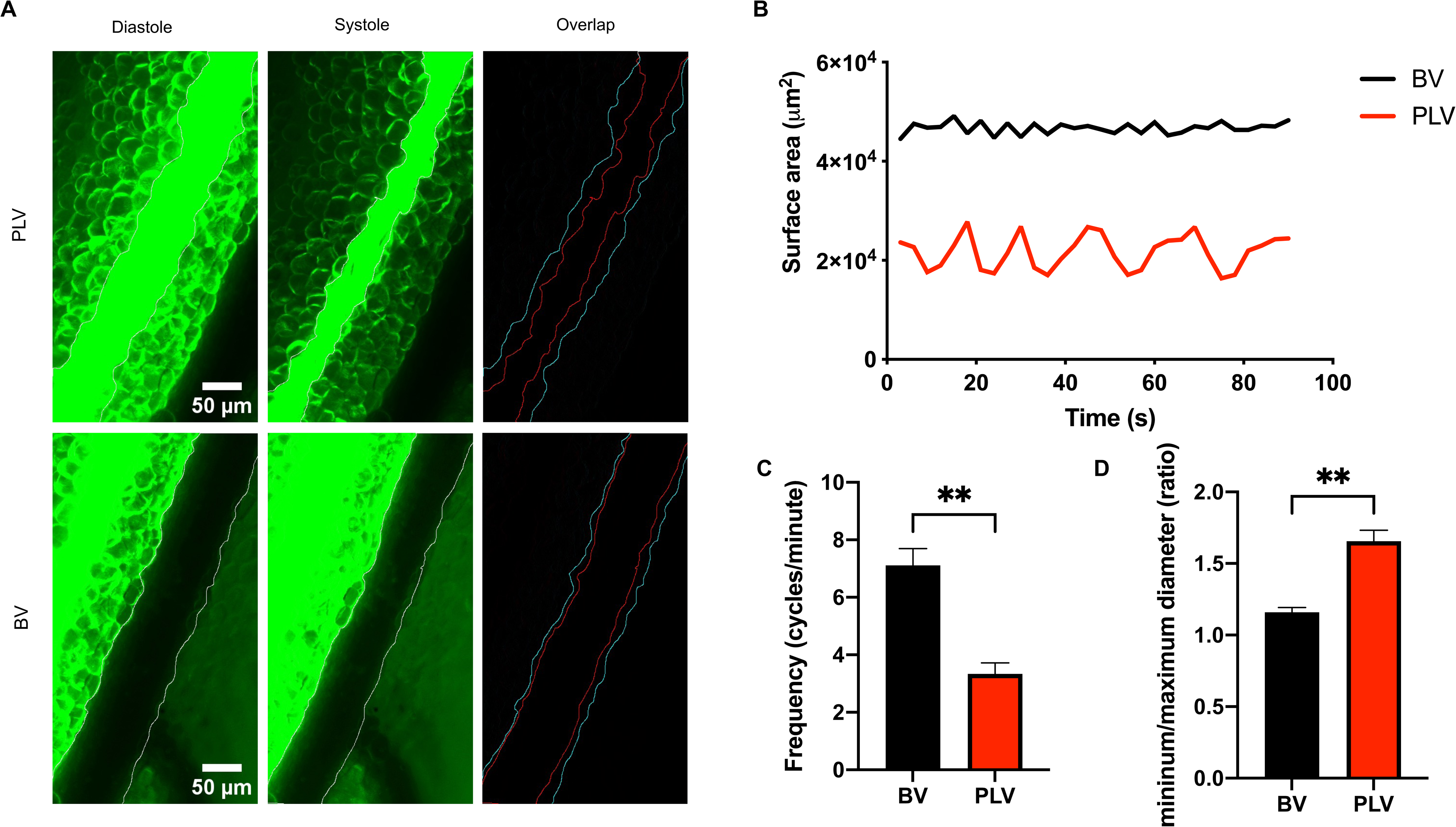
Popliteal lymphatic vessels and saphenous veins exhibit phasic contractility. **A.** Images of FITC-Dextran filled popliteal lymphatic vessels (PLVs) and adjacent saphenous veins (BVs). Diastole and systole images were taken at 45 and 57 seconds for PLVs and at 21 and 27 seconds for BVs. Diastole outline is cyan and systole outline is red. **B.** Representative area variation in PLVs and BVs over time. **C.** Frequency comparison between BV and PLV (n=3 mice). P-value calculated using unpaired student’s t-test **p<0.01. **D.** The maximum diastolic area and minimum systolic area for each vessel was calculated for each vessel and expressed as a ratio (n=3 mice). P-value calculated using student’s t-test **p<0.01.

### LMCs in popliteal lymphatic vessels and mesenteric lymphatic vessels do not derive from mature SMCs

Considering the functional and transcriptional similarities between LMCs and venous SMCs, we questioned whether saphenous vein SMCs might migrate to the popliteal lymphatic vessel and interconvert to form LMCs. This hypothesis is particularly intriguing given that blood vessels develop and are muscularized long before the formation of popliteal lymphatic vessels and subsequent investment of LMCs. To inform our lineage tracing approaches, we tracked the recruitment of LMCs to popliteal lymphatic vessels and mesenteric lymphatic vessels in embryos and postnatal mice by staining the hindlimbs and mesenteries, respectively with α-SMA (Fig. 4A-D). Prior studies show that LMCs appear by E18.5^2,10^. Indeed, E17.5 popliteal lymphatic vessels and mesenteric lymphatic vessels showed lymphatic vessels with LMC coverage. Following the emergence of LMCs on collecting LVs in the hindlimb and mesentery by E17.5, progressive expansion of LMCs occurs in early postnatal development as measured on P1 and P7 mice (Fig. 4E,F).

**Figure 4.**
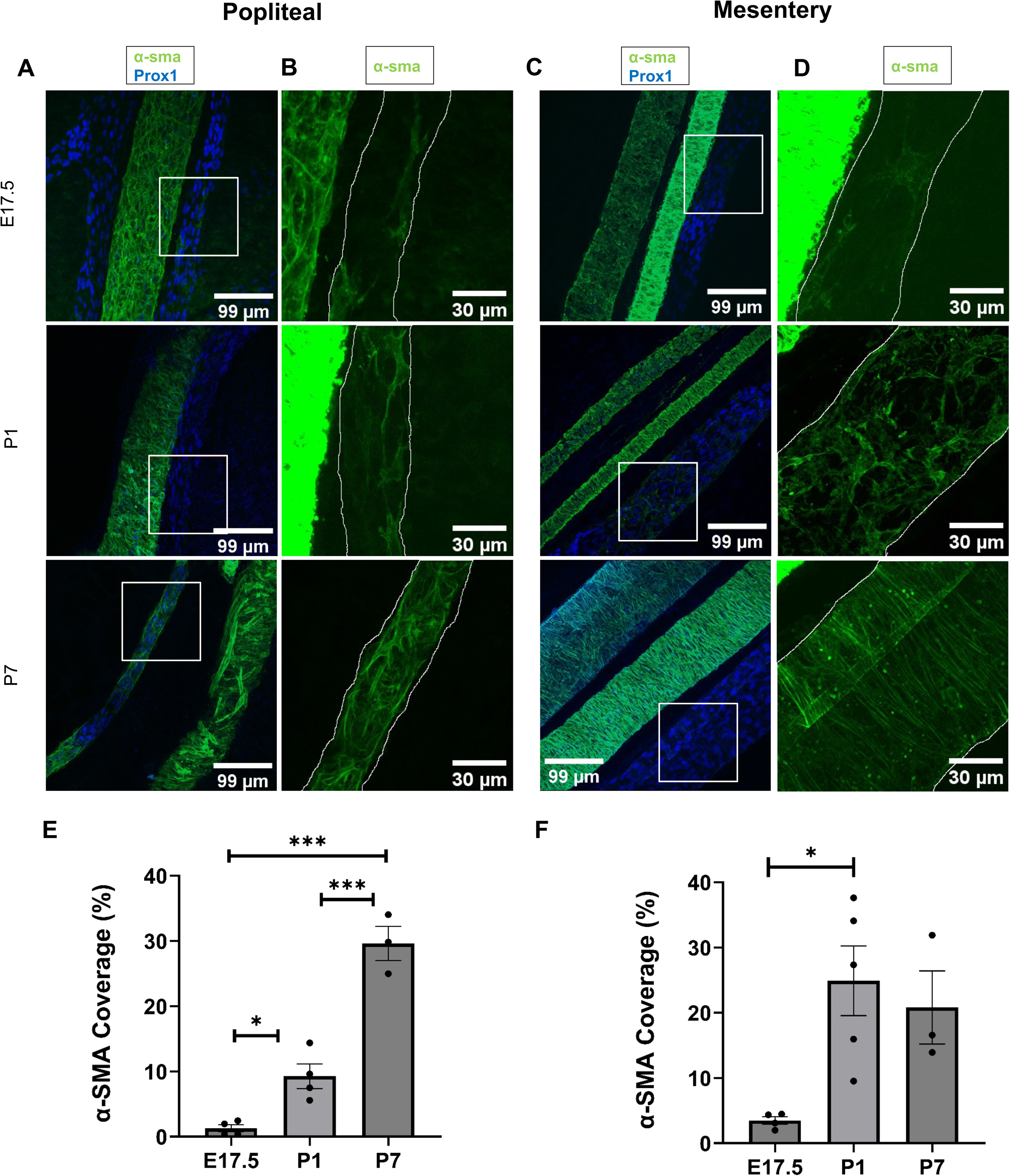
Time course of LMC appearance in hindlimb and mesentery. **A.** 3D projections of immunostained popliteal lymphatic vessels (PLV, squared) and adjacent saphenous vein of E17.5, P1 and P7 WT mice. Original magnification 40X; Scale bar 99 µm. **B.** Magnified areas of A. PLVs have been outlined. Scale bar 30 µm. **C.** 3D projections of immunostained mesenteric lymphatic vessels (MLV, squared) and adjacent blood vessels of E17.5, P1 and P7 WT mice. Original magnification 40X; Scale bar 99 µm. **D.** Magnified areas of B. MLVs have been outlined. Scale bar 30 µm. **E.** Percentage of α-sma covered area (mean and SEM) in PLVs of E17.5 (n=4), P1 (n=4) and P7 (n=3) WT mice. P-value calculated using ANOVA and Games-Howell post-hoc test. *p<0.05,**p<0.01, **p<0.001. **F.** Percentage of α-sma covered area (mean and SEM) in MLVs of E17.5 (n=4), P1 (n=5) and P7 (n=3) WT mice. P-value calculated using ANOVA and Games-Howell post-hoc test. *p<0.05.

Next, we performed lineage tracing using Myh11, a well-established marker of SMCs and LMCs^48,73^, to determine whether LMCs derived from Myh11+ SMCs. To achieve this, we crossed Myh11^CreERT2^ mice^28^ with Ai9-tdTomato reporter mice to generate Myh11 reporter mice that express tdTomato (tdT) upon cre recombination. Muscularization of blood vessels occurs at E10.5^74^, before the appearance of LMCs at E17.5. The development of mesenteric lymphatic vessels initiates around E14.5–15.5^11^. After confirming the lack of LMC coverage in mesenteric lymphatic vessels at E15.5 and at E17.0 (Fig.S2), at E14.0 we administered tamoxifen to pregnant dams to induce Cre expression and tdT recombination in embryos (Fig. 5A). Animals were harvested at P0, and as expected, arteries and veins in both hindlimbs and mesenteries showed coverage of tdT+ SMCs (in veins 0.36 and 0.03 positive cells per 100 µm^2^ respectively, p=0.16), but popliteal lymphatic vessels and mesenteric lymphatic vessels lacked coverage of tdT+ LMCs, despite exhibiting coverage of α-SMA+ LMCs (Fig. 5B-G). As a control, α-SMA+ LMCs of adult Myh11 reporter mice treated with tamoxifen (4 days before harvest) expressed tdT (Fig.S3). The absence of tdT+ LMCs at P0 suggests LMCs do not derive from mature SMCs or other cells expressing Myh11 at E14.0, and that LMCs may originate from immature progenitors, ultimately adopting a comparable expression profile as venous SMCs.

**Figure 5.**
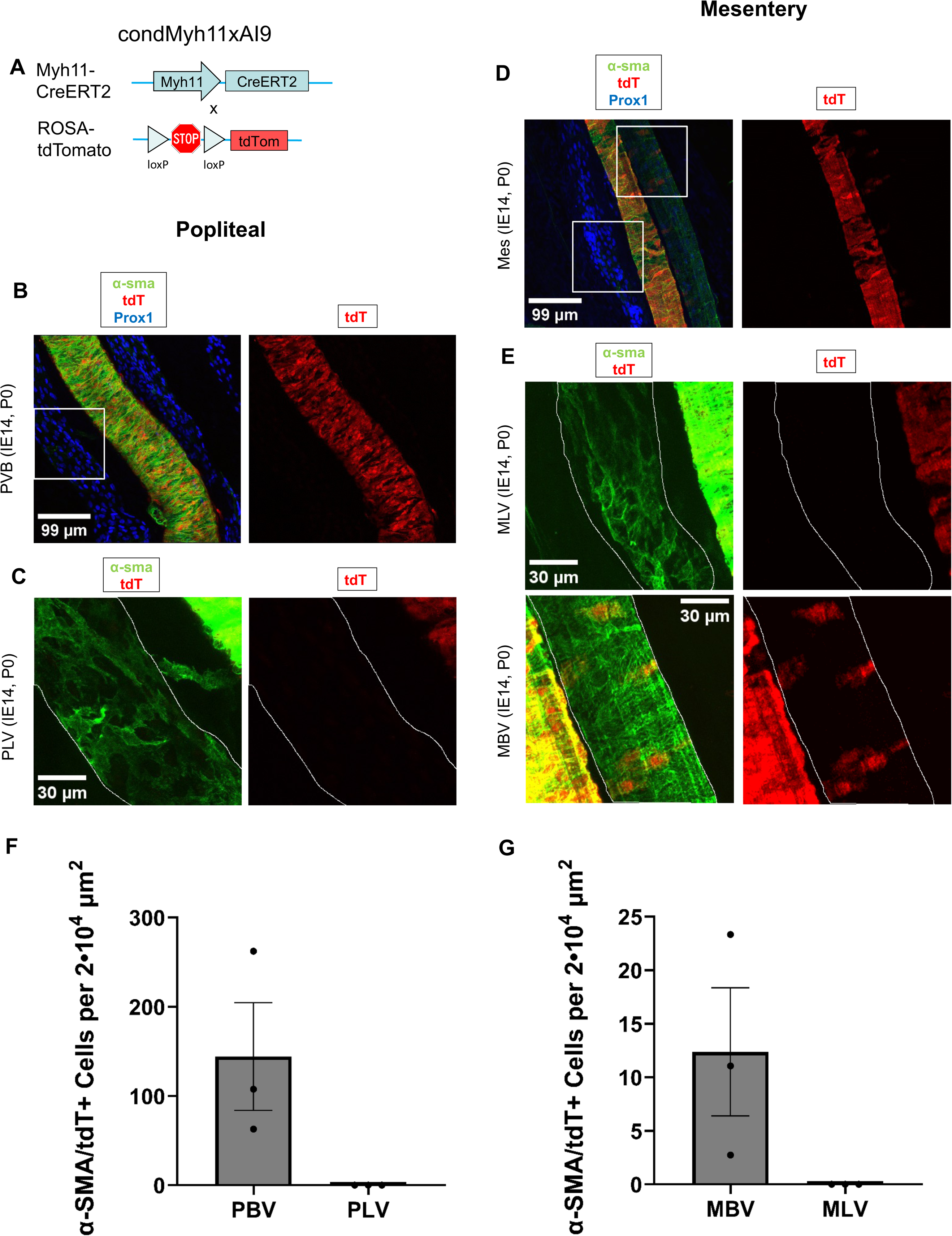
Lineage tracing reveals that LMCs are not derived from Myh11 expressing progenitors. **A**, Schematic of the recombination at the Rosa26 locus of Myh11CreERT2, Ai9-tdTomato mice after tamoxifen induction. **B.** 3D projections of immunostained popliteal vascular branch (PVB) consisting of popliteal lymphatic vessels (PLV, left one squared) and adjacent saphenous vein of P0 Myh11CreERT2, Ai9-tdTomato mice induced at E14. **C.** Magnified area of B. PLV has been outlined. **D.** 3D projections of immunostained mesenteric lymphatic vessels (MLV, left square) and adjacent blood vessel (artery in the center and vein on the right and squared) of P0 Myh11CreERT2, Ai9-tdTomato mice induced at E14. **E.** Magnified areas of D. MLV and vein (MBV) have been outlined. **F.** Counts of α-sma/tdTomato positive cells (mean and SEM) per 2·10^4^ µm^2^ in popliteal blood (PBV, n=3) and lymphatic (PLV, n=3) vessels. **G.** Counts of α-sma/tdTomato positive cells per 2·10^4^ µm^2^ in mesenteric vein (MBV, n=3) and lymphatic (MLV, n=3) vessels.

### Populations of LMCs and SMCs are derived from select mesodermal progenitors, but not from cardiomyocyte progenitors

Since SMCs were not a source of LMCs during development, we investigated whether both cell types originated from common progenitors. Because vascular SMCs have a primary derivation from the mesoderm^24^, we next asked whether the mesoderm is also a source of LMCs. To this end, we performed lineage tracing using WT1 (Wilms’ tumor suppressor gene), a marker of lateral plate mesoderm that has been shown to contribute to SMCs^27,75^. We crossed WT1^Cre^ mice with Ai9-tdT reporter mice to generate WT1 reporter mice (Fig. 6A).

**Figure 6.**
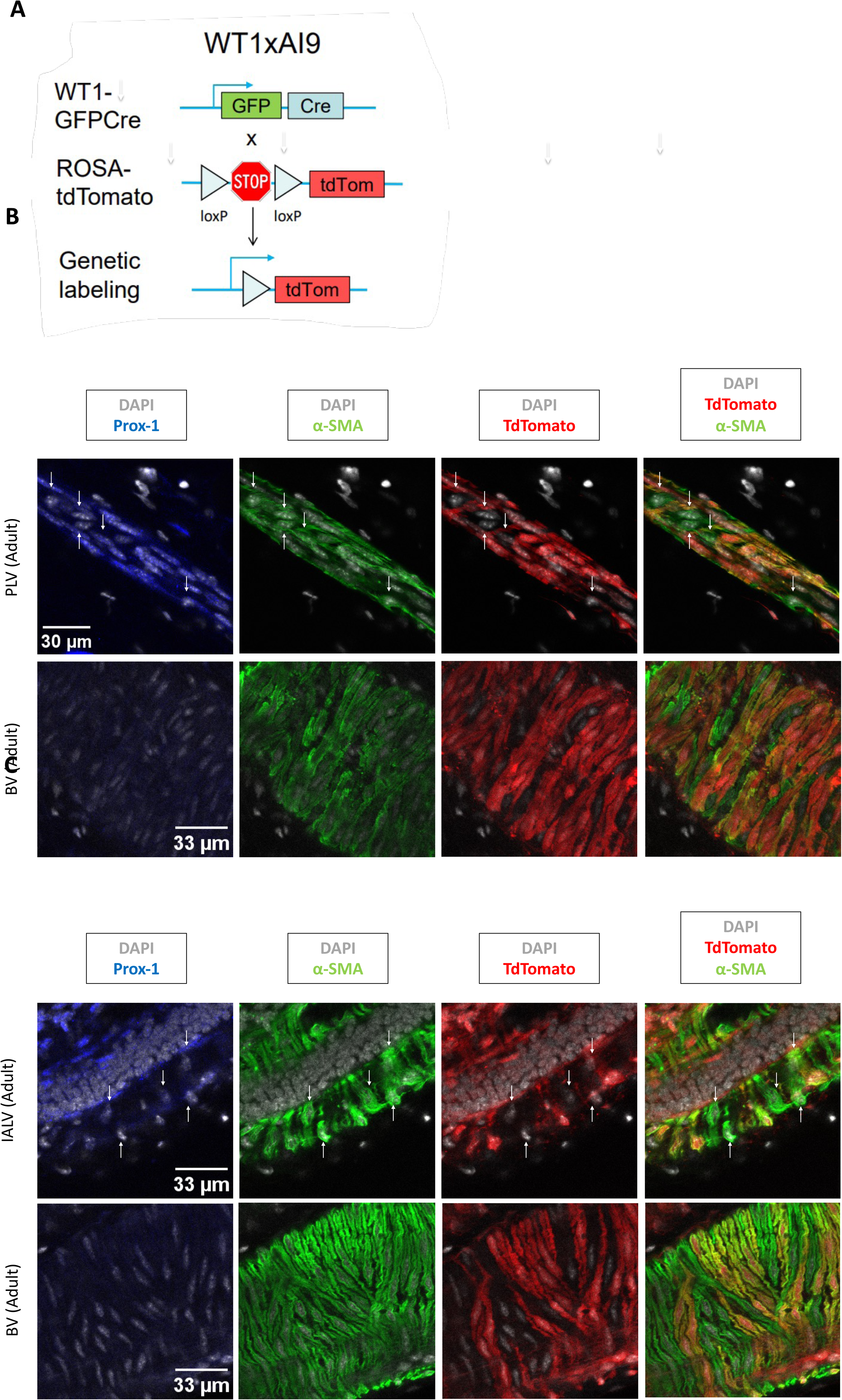
Lineage tracing reveals the contribution of WT1-expressing mesodermal progenitors to LMC and SMC populations in PLVs and IALVs. A, Schematic of the recombination at the Rosa26 locus of WT1GFPCre, Ai9-tdTomato mice. B, Single slice of Z-stack of immunostained popliteal lymphatic vessel (PLV) and saphenous vein (BV) of adult mice. Original magnification 40X; Scale bar 33 µm. Arrows point to α-SMA/tdTomato positive cells. C, Single slice of Z-stack of immunostained Inguinal-Axillary lymphatic vessel (IALV) and saphenous vein (BV) of adult mice. Original magnification 40X; Scale bar 33 µm. Arrows point to α-SMA/tdTomato positive cells. D, Counts (mean and SEM) of α-SMA/tdTomato positive lymphatic muscle cells (MCs) in adults (n=3) IALVs and PLVs.

We then isolated lymphatic vessels and adjacent blood vessels from the hindlimbs and the area extending from the inguinal region to the axillary region. Confocal imaging of whole mount-stained hindlimbs from neonatal and adult WT1 reporter mice revealed tdT+, α-SMA+ LMCs and SMCs associated with the popliteal lymphatic vessels and saphenous vein, respectively (Fig. 6B, S4A). Similarly, we found tdT+ LMCs and SMCs on the inguinal-axillary lymphatic vessel (IALV) and thoracoepigastric vein adjacent to the IALV (Fig. 6C). We found that tdT was primarily expressed in cells comprising vascular structures in the hindlimbs and inguinal-axillary regions (Fig. S4B,C). However, in both tissues tdT positive and tdT negative LMCs and SMCs were identified, suggesting heterogeneity in their lineage derivation. This heterogeneity was consistent throughout development as shown by the analysis of α-SMA/tdT cells per unit area at P7 compared to adult mice (0.24 vs 0.27 LMCs/100 µm^2^, p=0.806; 0.84 vs 0.65 SMCs/100 µm^2^, p=0.754; Fig. S4D).

To assess whether tdT labelling of LMCs and SMCs was derived from postnatal WT1 expression, we performed lineage tracing experiments by crossing WT1CreERT2 mice with Ai9-tdT reporter mice. The resulting mice were administered tamoxifen at P4 and P7, and tissues were collected at P11. As expected, kidney podocytes, which maintain expression of WT1 throughout adulthood, exhibited reporter expression, indicating successful induction (Fig. S4E). However, no tdT-labeled cells were detected in the popliteal lymphatic vessels or saphenous veins from hindlimbs of pups and suggests that tdT labeling is the result of embryonic WT1 expression by LMC and SMC progenitors (Fig. S4F).

To further assess the contribution of WT1+ progenitors to LMCs invested in popliteal lymphatic vessels, we used a reporter-independent mouse model. To this end, we created WT1 x DTR (diphtheria toxin receptor) mice by crossing WT1^Cre^ mice with ROSA26^iDTR^ mice. In the WT1x DTR mice, all cells that derive from WT1+ progenitors express DTR that renders them susceptible to diphtheria toxin-mediated ablation (Fig. 7A). To locally ablate hindlimb cells derived from WT1 expressing progenitors, adult mice were subjected to three consecutive days of diphtheria toxin (DT) subcutaneous injection into the hindlimb, and tissues were collected four days after the final DT application (day 7, Fig. 7B). Littermates negative for the WT1 transgene mice served as controls and were also administered DT. Whole mount staining of hindlimb tissue showed that DT administration reduced SMC coverage on the saphenous veins of WT1(Cre+) x DTR mice compared to WT1(Cre-) x DTR control mice (58% vs 34%, p=0.065, Fig. 7C,D). Further, popliteal lymphatic vessels from WT1(Cre+) x DTR mice had significantly lower LMC coverage compared to those of WT1(Cre-) x DTR control mice (63% vs 43%, p=0.006, Fig. 7E,F). These data corroborate findings from the WT1 reporter mice, indicating that a subset of LMCs and SMCs derive from WT1+ progenitors.

**Figure 7.**
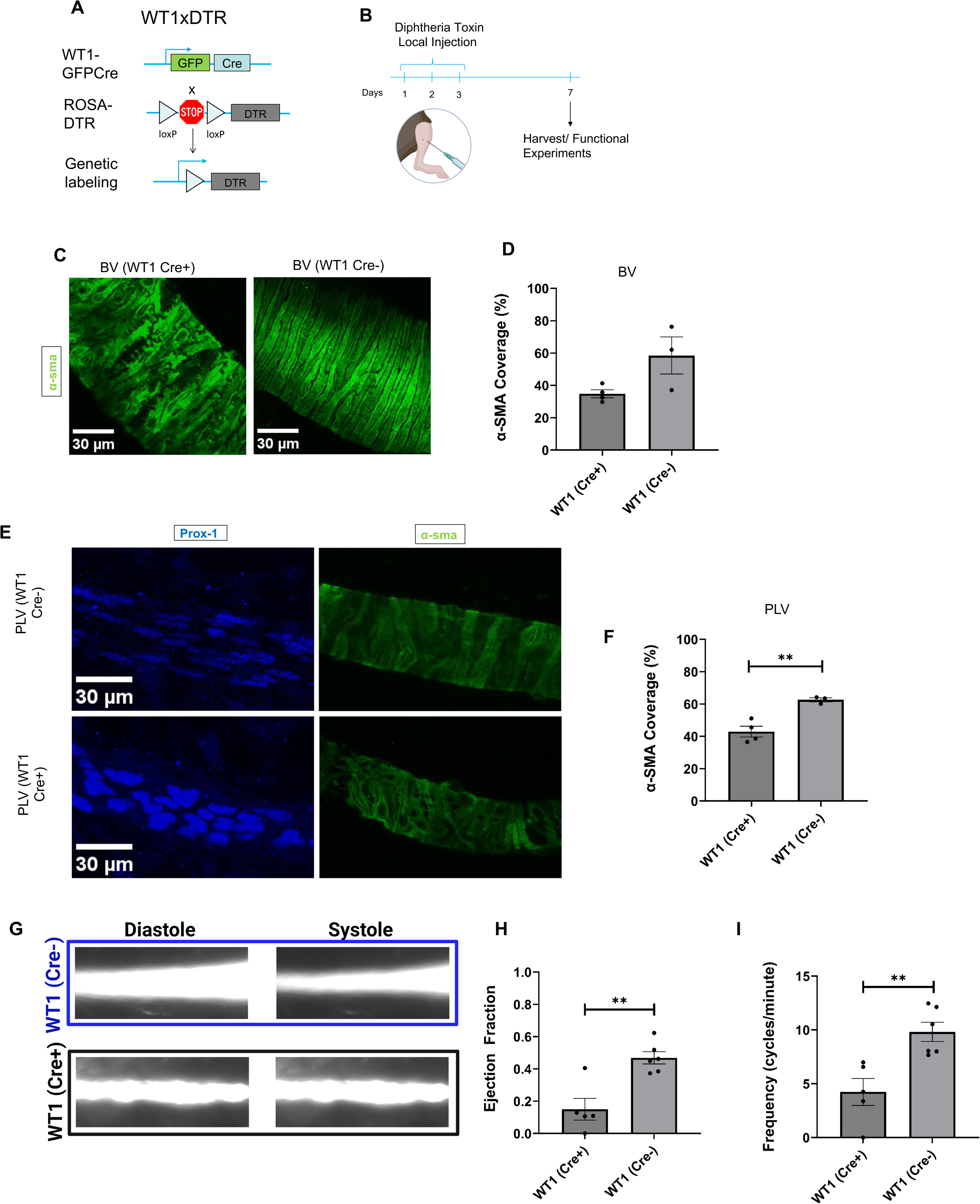
Ablation of WT1+ progenitor-derived LMCs attenuates SMC and LMC coverage of blood and lymphatic vessels, respectively. **A.** Schematic of the recombination at the Rosa26 locus of WT1GFPcre, Rosa-DTR mice. **B.** Timeline of diphtheria toxin treatment of WT1GFPcre, Rosa-DTR mice and controls (WT1Cre-). Cartoon created with Biorender. **C.** 3D projection of immunostained saphenous veins (BV) of WT1Cre+ and WT1Cre-adult mice. **D.** Percentage of α-sma covered area (mean and SEM) in BVs of WT1Cre+ (n=4) and WT1Cre-(n=3) adult mice. **E.** 3D projection of immunostained popliteal lymphatic vessels (PLVs) of WT1Cre+ and WT1Cre-adult mice. **F.** Percentage of α-sma covered area (mean and SEM) in PLVs of WT1Cre+ (n=4) and WT1Cre-(n=3) adult mice. P-value calculated by unpaired student’s T test. **p<0.01. **G.** FITC-Dextran filled PLVs of WT1Cre+ and WT1Cre-adult mice during diastole and systole. **H**. Ejection fraction (mean and SEM) in PLVs of WT1Cre+ (n=5) and WT1Cre-(n=6) adult mice. P-value calculated by Student’s T test. **p<0.01. **I.** Diameter variation (μm) over time (s) of PLVs of WT1Cre+ (n=5) and WT1Cre-(n=6) adult mice. P-value calculated by Student’s T test. **p<0.01

Next, we performed intravital imaging^34^ following DT application to measure whether WT1-derived cells were critical for the LMC-driven contraction of PLVs. The ejection fraction (EF) and frequency, indicators of lymphatic contractility, were significantly lower in the PLVs of WT1(Cre+) x DTR mice compared to controls (mean EF: 0.47 vs 0.15, p=0.002 and mean frequency: 4.2 vs 9.8, p=0.005, Fig. 7G-I). Together, these data suggest that WT1 progenitor-derived LMCs are necessary for lymphatic vessel contractility.

Finally, since LMCs have been reported to share functional and transcriptional characteristics with cardiomyocytes ^21,76^, we sought to test whether cardiac progenitors accounted for a population of tdT negative LMCs identified in WT1 reporter mice. To this end, we performed lineage tracing of Nkx2.5+ cardiac progenitors from the anterior lateral plate mesoderm. Nkx2.5 reporter mice were generated by crossing Nkx2.5^IRESCre^ mice with Ai9-tdT reporter mice (Fig. S5A). Confocal imaging of whole mount-stained hindlimbs revealed Nkx2.5-derived cells that localized near the saphenous vein, but no association of Nkx2.5-derived cells was observed in the popliteal lymphatic vessels (Fig. S5B). In addition to the hindlimbs, which contain the popliteal lymphatic vessels and adjacent blood vessels, we also harvested the mesenteries from P7 pups to assess whether the lack of contribution of Nkx2.5-expressing progenitors to LMCs was tissue-dependent. Similar to the hindlimb, Nkx2.5-expressing progenitors were observed in proximity to mesenteric blood vessels but were not associated with lymphatic vessels (Fig. S5C). Further, Nkx2.5-expressing progenitors that associated with blood vessels did not overlap with α-SMA positive SMCs.

## Discussion

We used scRNAseq to analyze the cellular composition and transcriptional profiles of cells obtained from murine popliteal lymphatic vessels and saphenous veins. From the hindlimb, we identified LMCs, venous SMCs, arteriole SMCs and weakly contractile perivascular cells^48^. Saphenous veins contained two subpopulations of SMCs and thus were more heterogeneous than LMCs of popliteal lymphatic vessels. The Adra2c-low subpopulation was enriched for similar genes as LMCs, implying comparable functional roles within the respective vessels. Adra2c-enriched SMCs expressed a distinct set of genes that were unique to this cell population. These cells likely have a specialized role in blood vessels, and given their enrichment of *P2rx1* and *Adra2c*, may be involved in neuronal regulation of blood vessel tone. Similarly, the role of the perivascular cell cluster is unknown. Although it is likely a contractile population because cells in this cluster express low levels of contractile genes, these cells may also provide structural support for vessels. Moreover, we cannot rule out the possibility that the perivascular cells are associated with venules embedded in the adipose surrounding the lymphatic vessels.

Despite many similarities in gene expression, we found differential expression of specific genes between LMCs and saphenous vein SMCs. The top differentially enriched genes are associated with signal transduction, muscle contraction, and ECM organization. In contrast, arteriole SMCs were notably distinguishable from LMCs and veSMCs. For example, arteriole SMCs expressed low levels of the canonical SMC markers *Actg2* and *Cnn1,* but were highly enriched in *Pln*, *Sorbs2,* and T-type channels, genes associated with cardiac contraction. Previous investigations have predominantly relied on bulk tissue analysis to study lymphatics and have implied smooth muscle and striated muscle genes expressed in LMCs. Because arteriole SMCs are enriched in proteins associated with cardiac contraction and intimately linked to lymphatic vessels, this may warrant a reevaluation of prior conclusions about the cardiac gene expression profile of LMCs. It is likely that arterioles closely associated with lymphatic vessels remain in single cell preparations due to the incomplete removal of the adipose tissue surrounding the vessels. Nevertheless, the reason for the enrichment of arteriolar smooth muscle cells relative to LMCs in our single-cell preparation through our digestion protocol remains unclear.

Given their transcriptional similarities, we tested whether LMCs are derived from SMCs by performing lineage tracing on Myh11 expressing cells in popliteal lymphatic vessels and mesenteric lymphatic vessels. The assessment of LMC labeling was made on P0 pups due to the high embryonic and neonatal mortality encountered with these animals after tamoxifen induction. While at this stage LMCs are still being added to the vessel, the number of LMCs was sufficient to conclude that they did not exhibit tdT labeling. In addition to suggesting that LMCs are not derived from SMCs, our results also show that neither LMCs or SMCs are derived from Myh11+ progenitors and that LMCs and SMC progenitors achieve terminal differentiation once associated with respective vessels.

To investigate whether LMCs and SMCs originate from a common progenitor, we directed our attention to the mesoderm. WT1 is a defining feature of the coelomic epithelium, a mesodermal structure that has been demonstrated to give rise to SMCs in numerous tissues^,27,75,77^. In the coelomic epithelium, WT1 is thought to facilitate the epithelial-mesenchymal transition (EMT) required to produce the mesenchymal progenitors that ultimately differentiate into various cell types^78,79^. WT1 also serves as a lateral plate mesoderm marker^78–80^ as it is expressed within the coelomic space of the lateral plate mesoderm^81^. Through lineage tracing with a conventional WT1 Cre driver, we show that WT1+ mesodermal progenitors are a source of LMCs in mouse popliteal lymphatic vessels and inguinal-axillary lymphatic vessels. Additionally, through lineage tracing with an inducible WT1 Cre driver we show that WT1 is not expressed in mature LMCs.

Although LMCs have been reported to share contractile proteins with cardiomyocytes, our study revealed that neither LMCs (or SMCs) from the hindlimb or mesentery were derived from Nkx2.5-expressing progenitors. From a developmental approach, this is consistent with the origin of both tissues since Nkx2.5+ progenitors are derived from the anterior lateral plate mesoderm, while both the hindlimbs and gut form mostly from the posterior lateral plate mesoderm^76^. Based on a recent study demonstrating that LMCs develop from pathways independent of skeletal muscle tissue^18^, it appears that neither cardiac (this study) or skeletal muscle progenitors confer striated muscle genes to LMCs.

Our finding that LMCs have a mesodermal origin aligns with prior research by Kenney *et al.* demonstrating the expression of a mesenchymal signature of LMC progenitors^18^. Notably, LMCs in our study maintained expression of Prrx1, indicating that this mesenchymal marker is maintained throughout adulthood in LMCs. Also, in agreement with the lineage tracing studies performed by Kenney *et al*^18^, *Cspg4*-expressing cells were not detected in adult LMCs. Our data show that *Cspg4* was enriched in arteriolar SMCs, which is also consistent with prior literature^48,61^. However, several of our findings contrast with scRNAseq results of LMCs and SMCs from popliteal lymphatic vessels^82^. In the study by Kenney *et al.*, the LMCs and SMCs from popliteal vessels accounted for a large percentage of the total cells, but neither the LMCs nor SMCs expressed the canonical markers of contractile cells. Instead, *Pi16* was one of the most enriched markers. In our study, *Pi16* was enriched in fibroblasts, but was not expressed by contractile cells. Indeed, a recent study shows that fibroblasts are a prominent cell type in the adventitia of lymphatic vessels^83^.

In conclusion, in this study we show that LMCs and venous SMCs express many common genes, suggesting that they exhibit a transcriptional program that exists on a continuum with varying levels of gene expression. These findings have therapeutic implications, as the similar characteristics between LMCs raise concerns about potential off-target effects when targeting one cell type. On the other hand, it also presents an opportunity to harness established therapies for SMCs to modulate LMC function. A limitation of the study is that the transcriptional analysis was limited to one tissue, and LMCs and SMCs likely display tissue-specific transcriptional heterogeneity. Although we uncovered the contribution of WT1+ mesodermal progenitors LMC and SMCs during embryonic development, an additional limitation of the study is that it is unclear which additional embryonic origins LMCs may arise from, since not all LMCs were derived WT1+ progenitors. Finally, we demonstrate that saphenous veins in the murine hindlimb exhibit phasic contractility. Saphenous veins had a higher contraction frequency than popliteal lymphatic vessels, but popliteal lymphatic vessels exhibited stronger contractions. These contractile differences may arise from the varying luminal pressures, diameters, and thickness^84^ between the vessels, alongside unique responses to various stimuli.

## Supporting information

SFigs1-5

## Acknowledgements

We thank the Single Cell Sequencing Core and Cellular Imaging Core of Boston University Chobanian & Avedisian School of Medicine. We also thank Katarina Ruscic and Johanna Rajotte from Massachusetts General Hospital and Harvard Medical School for assistance with contractility experiments.

