## Supplementary material for "Transcriptional, developmental, and functional parallels of lymphatic and venous smooth muscle": SFigs1-5

### Figure S1

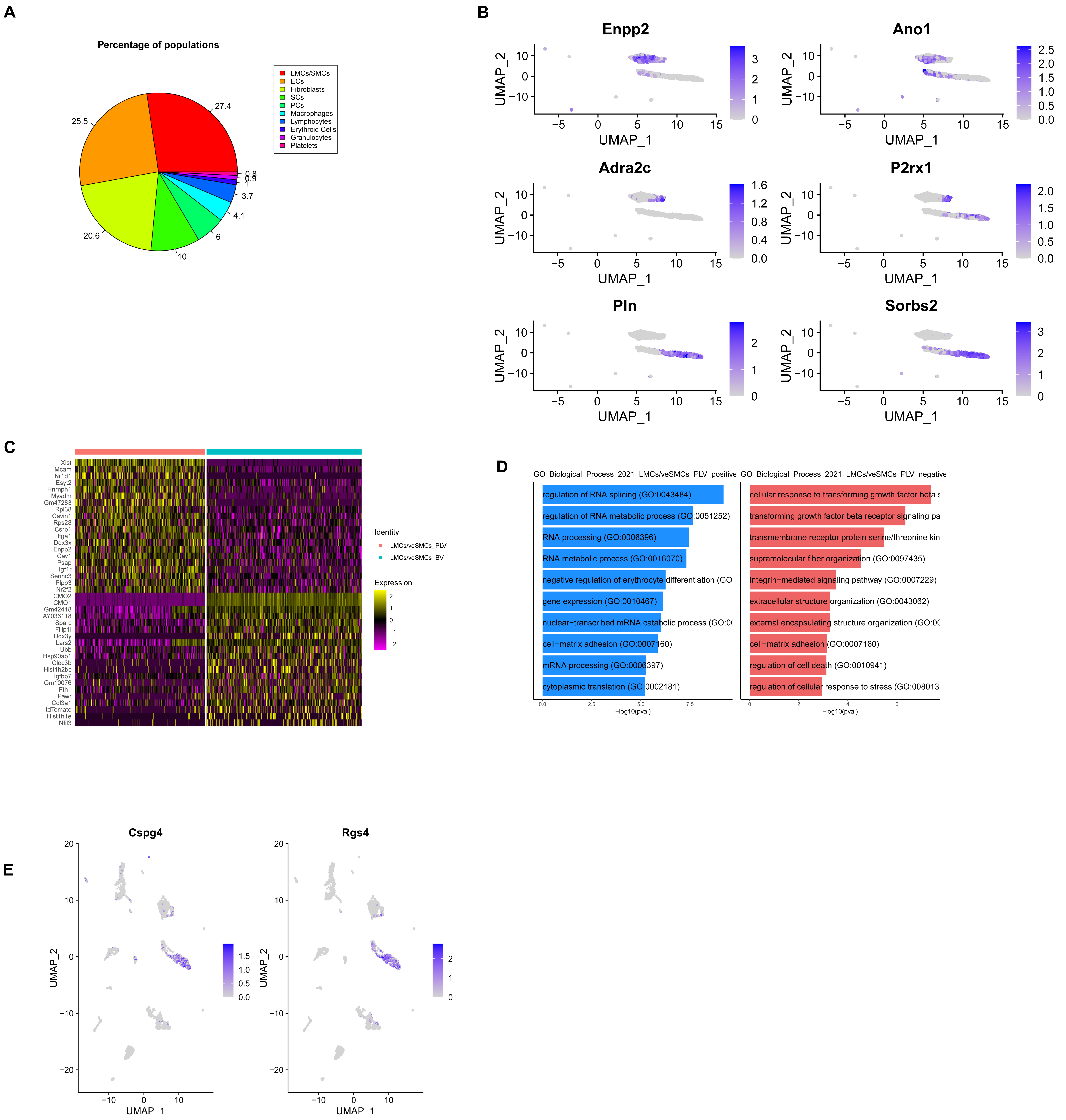


**Supplemental Figure 1. A.** Pie chart of the percentage of most relevant cell types and subtypes. **B.** UMAP visualization (gray, low; blue, high expression) of genes enriched in groups of the contractile clusters. **C.** Heatmap of the differentially expressed genes between LMCs and veSMCs. **D.** Enricher GO analysis (GO_Biological_Process_2021) of the differentially expressed genes between LMC and veSMC clusters. **E.** UMAP visualization (gray, low; blue, high expression) of genes enriched in aSMCs.

### Figure S2

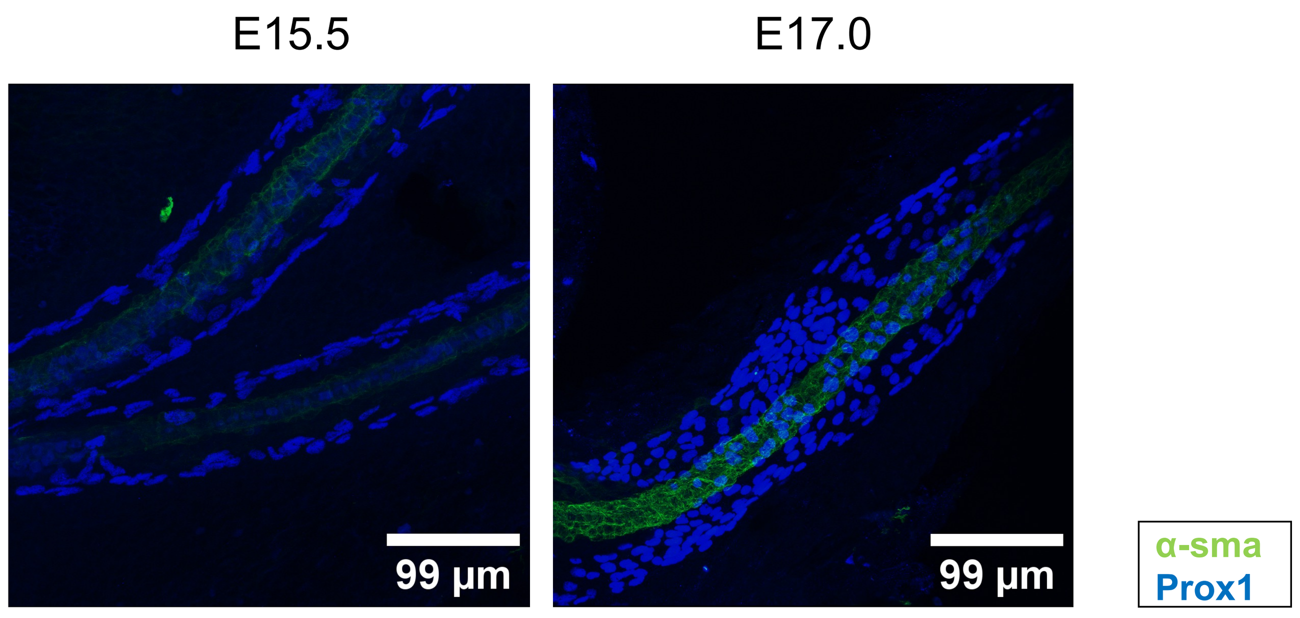


**Supplemental Figure 2.** 3D projections of immunostained mesenteric lymphatic vessels of E15.5 and E17.0 mouse embryos. Original magnification 40X; Scale bar 99 µm.

### Figure S3

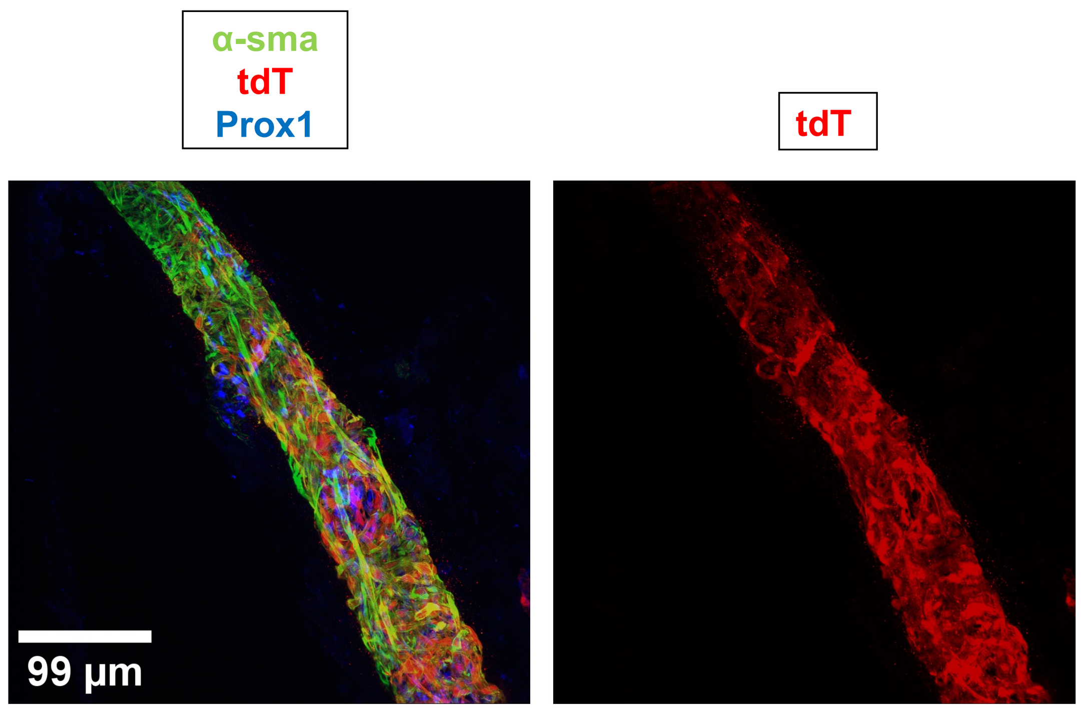


**Supplemental Figure 3.** 3D projection of immunostained popliteal lymphatic vessel of adult Myh11CreERT2, Ai9-tdTomato mice 4 days after tamoxifen induction. Original magnification 40X; Scale bar 99 µm.

### Figure S4

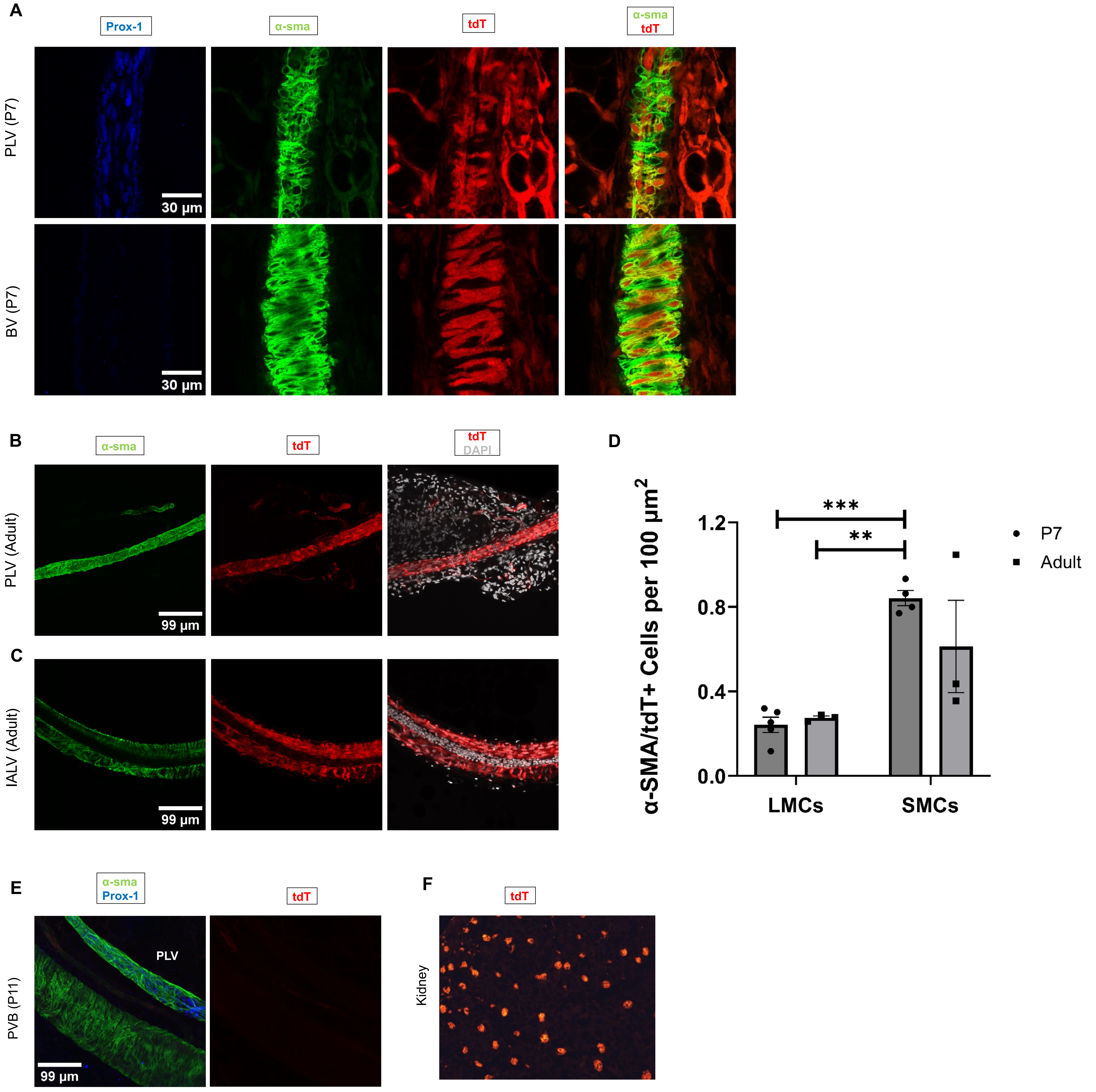


**Supplemental Figure 4. A.** Selected single slices of Z-stack of immunostained popliteal lymphatic vessel (PLV) and saphenous vein (BV) of P7 WT1 reporter mice. Prox1 panels show EC/LEC prominent stacks, α-sma and tdT panels show SMC/LMC prominent stacks. Original magnification 40X; Scale bar 30 μm. **B.** 3D projections of immunostained popliteal lymphatic vessel (PLV) of an adult WT1GFPCre, Ai9-tdTomato (WT1 reporter) mouse. **C.** Single slice of Z-stack of immunostained inguinal-axillary lymphatic vessel (IALV) of an adult WT1 reporter mouse. **D.** Counts (mean and SEM) of α-sma/tdTomato positive cells per 100 μm$^2$ in P7 (n=4 mice) and adult (n=3 mice) saphenous vein (SMCs) and popliteal lymphatic vessels (LMCs) of WT1 reporter mice. Area refers to surface within vessel outlines in 3D projection images (single measurement). P-value calculated using ANOVA and Games-Howell's post-hoc test. *** p<0.001, **p<0.01. **E.** 3D projection of immunostained popliteal vascular branches (PVB) of P11 WT1CreERT2, Ai9-tdTomato (conditional WT1) mice induced postnatally (P4,P7). Image shows a lymphatic vessel and the adjacent saphenous vein. **F.** Cryosections of kidneys of conditional WT1 mice induced at P4,P7.

### Figure S5

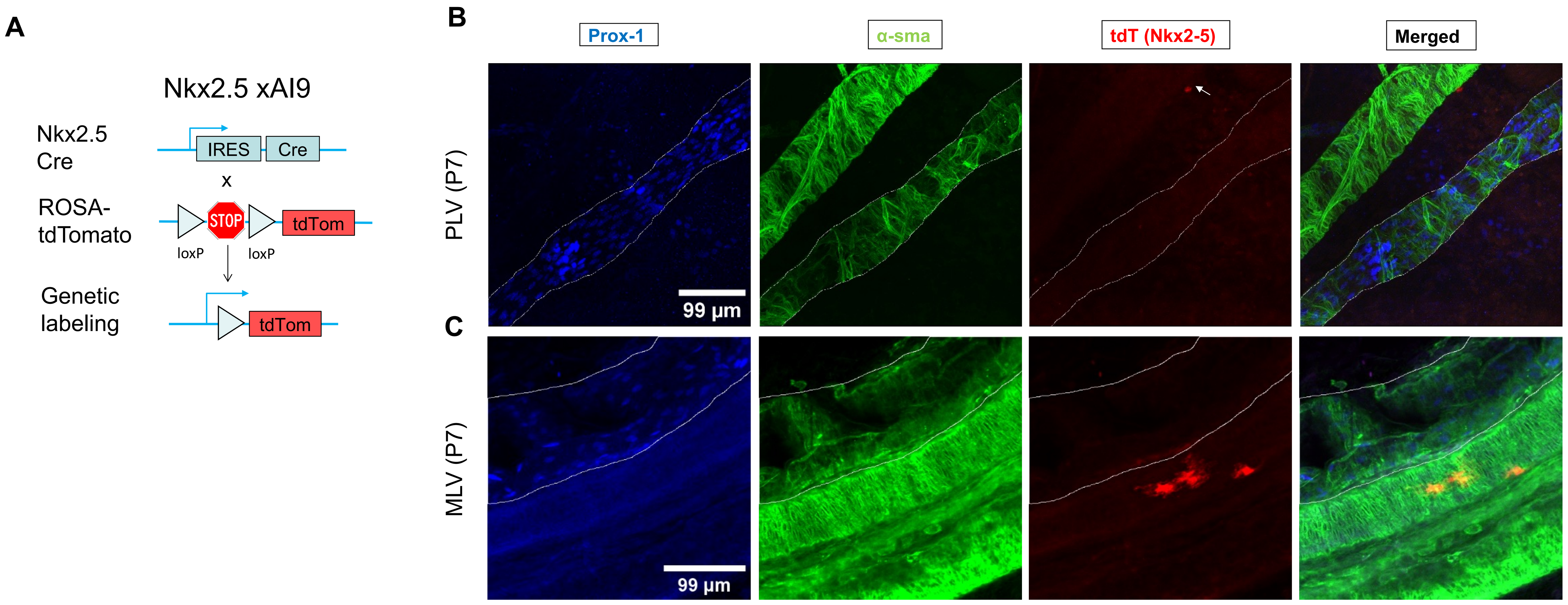


**Supplemental Figure 5. A.** Schematic of the recombination at the Rosa26 locus of Nkx2.5IRESCre, Ai9-tdTomato (Nkx2.5 reporter) mice. **B.** 3D projections of immunostained PLV (outlined) and adjacent saphenous vein of a P7 Nkx2.5 reporter mouse. Arrow points to tdTomato positive extravascular cell. Original magnification 40X; Scale bar 99 µm. **C.** Single slice of Z-stack of immunostained mesenteric lymphatic vessel (outlined) and adjacent blood vessels of P7 Nkx2.5 reporter mouse with extravascular tdTomato positive cells. Original magnification 40X; Scale bar 30 µm.
